# Older adults benefit from more widespread brain network integration during working memory

**DOI:** 10.1101/642447

**Authors:** C.A. Crowell, S.W. Davis, L. Beynel, L. Deng, D. Lakhlani, S.A. Hilbig, H. Palmer, A. Brito, J. Wang, A. V Peterchev, B. Luber, S.H. Lisanby, L.G. Appelbaum, R. Cabeza

## Abstract

Neuroimaging evidence suggests that the aging brain relies on a more distributed set of cortical regions than younger adults in order to maintain successful levels of performance during demanding cognitive tasks. However, it remains unclear how task demands give rise to this age-related expansion in cortical networks. To investigate this issue, we used functional magnetic resonance imaging to measure univariate activity, network connectivity, and cognitive performance in younger and older adults during a working memory (WM) task. In the WM task investigated, participants hold letters online (maintenance) while reordering them alphabetically (manipulation). WM load was titrated to obtain four individualized difficulty levels. Network integration—defined as the ratio of within-versus between-network connectivity—was linked to individual differences in WM capacity. The study yielded three main findings. First, as task difficulty increased, network integration decreased in younger adults, whereas it increased in older adults. Second, age-related increases in network integration were driven by increases in right hemispheric connectivity to both left and right cortical regions, a finding that helps to reconcile extant theories of compensatory recruitment in aging to address the multivariate dynamics of global network functioning. Lastly, older adults with higher WM capacity demonstrated higher levels of network integration in the most difficult condition. These results shed light on the mechanisms of age-related network reorganization by suggesting that changes in network connectivity may act as an adaptive form of compensation, with older adults recruiting a more distributed cortical network as task demands increase.

**Significance statement:** Older adults often activate brain regions not engaged by younger adults, but the circumstances under which this widespread network emerges are unclear. Here, we examined the effects of aging on network connectivity between task regions recruited during a working memory (WM) manipulation task, and the rest of the brain. We found an age-related increase in the more global network integration in older adults, and an association between this integration and working memory capacity in older adults. The findings are generally consistent with the compensatory interpretation of these effects.

## Introduction

Despite substantial anatomical and functional decline, the aging brain retains a surprising degree of neural plasticity. In functional neuroimaging studies, for example, older adults often activate brain regions not engaged by younger adults during the same tasks (Park and Reuter-Lorenz, 2009; Cabeza and Dennis, 2013). Although over-recruitment in older adults is often interpreted as compensatory (Cabeza et al., 2018), it is unclear if and how the regions over-recruited by older adults are integrated with the network mediating task performance. To investigate this question, the current study assessed the effects of aging on network integration, an established concept of interest in characterizing brain dynamics (Tononi et al., 1994), during a working memory (WM) manipulation task. The study had three main goals.

The first goal of the study was to examine how age effects on network integration differ as a function of WM demands. As the number of items maintained in WM (load) increases, brain activity tends to rise monotonically in several regions, including dorsolateral prefrontal cortex (DLPFC) and lateral parietal cortex (LPC, Braver et al., 1997; Rypma et al., 1999; Beauchamp et al., 2001; Veltman et al., 2003). These activations tend to increase more rapidly in older than younger adults up to a certain level of WM demands (Low et al., 2009; Cappell et al., 2010; Schneider-Garces et al., 2010). According to the Compensation-Related Utilization of Neural Circuits hypothesis (CRUNCH, Reuter-Lorenz and Cappell, 2008), due to processing inefficiencies, older adults over-recruit neural resources at lower levels of task difficulty than younger adults. What is unclear from available CRUNCH evidence is whether the accelerated brain recruitment in older adults is limited to the activation of individual regions or whether it involves a reorganization of the underlying task network. We hypothesized that, as task demands increase, the WM network would become more integrated with the rest of the brain and this effect would be greater for older than younger adults (Hypothesis 1).

The second goal of this study was to examine hemispheric differences in network integration for young and older adults. In functional neuroimaging studies, activations are often more bilateral in older adults than younger adults, an effect known as Hemispheric Asymmetry Reduction in Older Adults (HAROLD, Cabeza, 2002). For example, during a verbal WM task that yields left lateralized activations in younger adults, older adults may show additional activity in the right hemisphere (Reuter-Lorenz et al., 2000). As in the case of CRUNCH, most evidence for HAROLD is based on univariate activity in individual regions, and hence, the network mechanisms of age-related hemispheric differences remain uncertain. We hypothesized that during the left-lateralized verbal WM task, older adults would show greater demand-related network integration in the right-hemisphere (Hypothesis 2).

The third goal of the study was to investigate if age-related changes in network integration relate to individual differences in WM performance. It has been suggested that more widespread activity in older adults is beneficial for performance (Park and Reuter-Lorenz, 2009; Cabeza and Dennis, 2012; Cabeza and Dennis, 2013), and both CRUNCH and HAROLD effects have been interpreted as compensatory. However, the evidence for compensation has been mostly based on univariate activity and evidence that network changes in older adults contribute to cognitive performance is largely missing (however, see Monge et al., 2018). We hypothesized that age-related WM network integration would be associated with WM ability in older adults (Hypothesis 3).

To test these hypotheses, participants completed a verbal WM manipulation task in which they maintained consonants briefly in memory while mentally rearranging them into alphabetical order. fMRI analyses focused on the effects of WM load on delay-period functional connectivity, while the WM network was defined as regions activated by the task irrespective of load. Here, network integration was measured as the ratio of within-versus between-network connectivity, while network integration was compared within and between hemispheres, and correlated with WM capacity to test the three stated hypotheses.

## Materials and Methods

### Participants

Forty-four young adults aged 18 to 35 (mean 22.8 ± 4.6) and 32 older adults aged 60 to 80 (mean 69.1 ± 5.3) participated in the study for monetary compensation and consented to the protocol approved by the Duke Medical School IRB (#Pro00065334). These participants were enrolled in a 6-day TMS protocol (Beynel et al., 2019; Beynel et al., in review), but only data from the screening session (Day 1) and MRI session (Day 2) are reported here. Participants had no history of psychiatric or neurological disorders and were not using psychoactive drugs. Participants were excluded because of poor functional imaging quality (excessive movement or falling asleep during data acquisition, n=3), or due to poor task performance in the scanner (accuracy greater than two standard deviations below the group mean, n=6). Thus, 37 young adults and 30 older adults were included in the final analysis.

### Experimental Design and Statistical Analyses

#### Behavioral procedure and task design

Participants performed a verbal *WM manipulation task* (**Fig. 1A**). In this task, an array of 3 to 9 consonant letters was presented for 3 seconds followed by a 5-second delay period, during which participants mentally rearranged letters into alphabetical order. Vowels were excluded to prevent chunking. After the delay period, a letter and number were presented together for 4 seconds and the participants pressed one of three buttons to indicate if the probe letter (1) appeared in the position indicated by the number in the alphabetized list *(Valid,* 40% of trials), (2) was part of original set but the number did not match the position in the alphabetized list *(Invalid,* 40% of trials), or (3) was not part of the original set *(New,* 20% of trials). These three types of trials occurred in random order. During the subject-specific titration on Day 1 (see the following paragraph), the response phase was followed by a 5-second inter-trial interval (ITI). During practice, participants were given feedback during this ITI on their accuracy after each trial, and at the end of each block. Twenty-five trials were included in each of the 6 blocks with a brief, self-paced rest interval between blocks.

**Figure 1.**
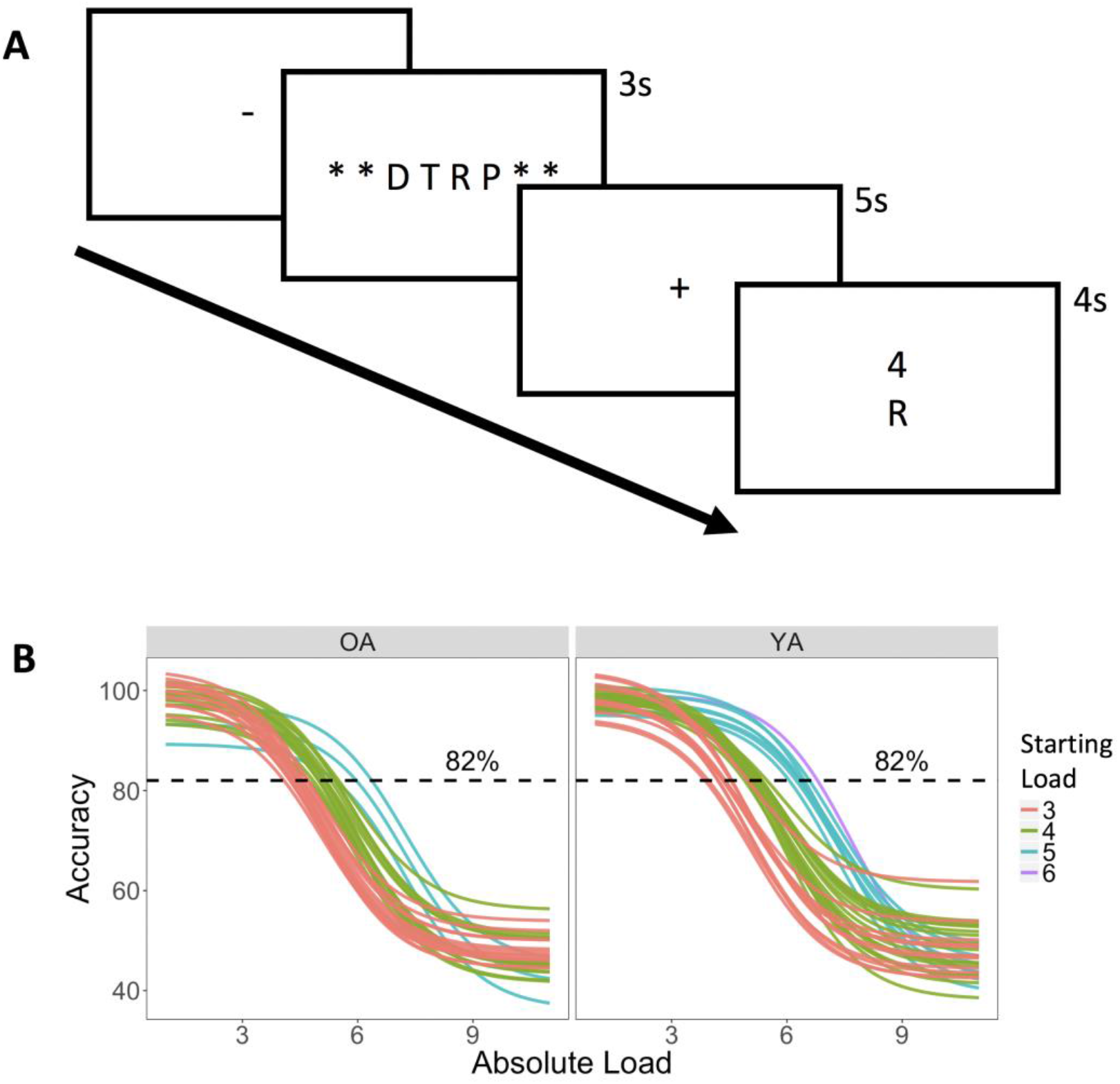
Task paradigm and individual WM load determination. **(A)** Illustration of WM manipulation during an array of 3-9 letters was presented, followed by a delay during which participants were asked to report if a probe number matches the serial position of a probe letter, once the original array is rearranged in alphabetical order. **(B)** Individually-titrated Difficulty Levels were determined using sigmoid curves fitted to individual performance data from screening visit, based on accuracy.

As part of the overall protocol, subjects participated in up to 6 experimental sessions, but only the first two are relevant to this study. In the first session, participants performed the WM manipulation task outside the scanner in order to identify the range of WM loads producing parametric changes in performance for each participant. The optimal load was identified using 2-down-1-up staircase procedure: when a trial was answered correctly, the load was increased by 1, and when it was answered incorrectly, the load was decreased by 2. New trials were excluded from the analysis and accuracy data, collapsed across Valid and Invalid trials at each load, were then fitted to a sigmoid function with performance threshold criterion set at 82% accuracy. To ensure that the psychometric function was not strongly influenced by noise in loads with a low number of trials, 50% accuracy was used for the largest loads if less than 10 trials were tested. To achieve more stable curve fits, peripheral anchors were added by including points for loads of 1 and 2 at 100% accuracy and loads 10 and 11 at 50% accuracy. Four individualized difficulty levels were defined according to the intersection between a sigmoid curve, fit to the data, and an 82% accuracy threshold (the criterion). The two loads below this intersection were defined as the Very Easy and Easy levels, and the two loads above were defined as the Medium and Hard levels. Thus, the four Absolute Loads selected for an individual depended on his/her WM ability (e.g., 3-4-5-6 letters in one participant, 4-5-6-7 in another participant). As such, for clarity the Very Easy, Easy Medium and Hard conditions are referred to as *WM Difficulty Levels.* Individual sigmoid curves for all young and older adults, and subsequent determination of their Starting Load (i.e. the Absolute Load value of the Very Easy condition), are shown in **Figure 1B**.

In the second session, participants performed the WM manipulation task inside the MRI scanner. Four blocks, each with 30 trials, were performed using the 4 individually-titrated Difficulty Levels. Stimuli were back-projected onto a screen located at the foot of the MRI bed using an LCD projector. Subjects viewed the screen via a mirror system located in the head coil and the start of each run was synchronized with the MRI acquisition computer. Trial-by-trial feedback was not given, but the overall block accuracy was presented at the end of each block. Behavioral responses were recorded with a 4-key fiber-optic response box (Resonance Technology, Inc.). Scanner noise was reduced with ear plugs, and head motion was minimized with foam pads. When necessary, vision was corrected using MRI-compatible lenses that matched the distance prescription used by the participant. The total scan time, including breaks and structural scans, was approximately 1 hour 40 minutes.

#### MRI scanning and data preprocessing

MRI was performed in a 3-T GE scanner at the at Duke Brain Imaging Analysis Center (BIAC). Structural MRI and diffusion-weighted imaging (DWI) scans were followed by performing 4 fMRI runs of the WM manipulation task. The anatomical MRI was acquired using a 3D T1-weighted echo-planar sequence (matrix = 256 x 256, time repetition (TR) = 7.15 ms, time echo (TE) = 2.7 ms, field of view (FOV) = 256mm^2^, slices = 196, slice thickness = 1 mm). In the fMRI runs, coplanar functional images were acquired using an inverse spiral sequence (64 × 64 matrix, TR = 2000 ms, TE = 25 ms, FOV = 220mm^2^, 34 slices, 4 mm slice thickness, 280 images). Finally, DWI data were collected using a singleshot echo-planar imaging sequence (TR = 17000 ms, slices = 76, thickness = 2.0 mm, FOV = 256 × 256 mm^2^, matrix size 128 × 128, voxel size = 2 mm^3^, b value = 2000 s/mm^2^, diffusion-sensitizing directions = 26, total images = 960, total scan time = 7.5 min).

Functional images were preprocessed using image processing tools, including FLIRT and FEAT from the fMRIB Software Library (FSL), in a publicly available pipeline developed by the Duke Brain Imaging and Analysis Center (https://wiki.biac.duke.edu/biac:analysis:resting_pipeline). Images were corrected for slice acquisition timing, motion, and linear trend; motion correction was performed using MCFLIRT, and 6 motion parameters estimated from the step were then regressed out of each functional voxel using standard linear regression. Images were then temporally smoothed with a high-pass filter using a 190-second cutoff and normalized to the Montreal Neurological Institute (MNI) stereotaxic space. White matter (WhM) and CSF signals were also removed from the data, using WhM/CSF masks generated by FSL’s FAST and regressed from the functional data using the same method as the motion parameters. Spatial filtering with a Gaussian kernel of full-width half-maximum (FWHM) of 8 mm was applied.

#### Behavioral analyses

Accuracy and reaction times (RTs) of WM manipulation trials were analyzed for each individually-titrated WM Difficulty Level. RTs were analyzed, only for correct trials (74.6% of total trials), using a linear restricted maximum likelihood model. Accuracy was analyzed using a binomial logistic regression model including all trials. In both models, R (R Core Team, 2012) and lme4 (Bates et al., 2012) were used to perform mixed effects analysis. WM Difficulty Level was entered as a fixed effect, and both model intercepts and by-subject random slopes across WM Difficulty Levels were included as random effects. Gender and each subject’s WM capacity were also included to account for standardizing difficulty levels across subjects and general differences in task ability. No deviations from homoscedasticity or normality were observed. P-values were obtained by likelihood ratio tests of the full model with the variable in question against a null model without the variable in question using an ANOVA to compare the model fits. There was no missing data, but in 1.8% of trials participants failed to respond within the 4-second response window (143 out of 8040 trials); these trials were excluded from all analyses.

#### fMRI analyses

A parametric approach was used to investigate how activity varied as a function of WM load. First-level voxel time-series analysis was carried out using general linear modeling (GLM); fixed effects models were carried out to examine the effects of load; separate events were modeled for the array presentation (duration: 3 seconds), delay period (duration: 5 seconds), and response (duration: subject reaction time), each with an onset at the beginning of the event. The delay period was additionally split into four separate regressors to model each of the WM Difficulty Levels for each subject. Incorrect and non-response trials were modeled identically, but separately, and were not considered in the univariate results below. Subsequent to individual-level models, random-effects analysis was performed on parameter estimates of the delay-period regressors (p < 0.005, cluster correction: z > 2.0). Subsequent group-level analyses were performed across the four delay-period regressors in SPM12.

#### Cortical parcellation

Before functional matrices were constructed, a consistent parcellation scheme was established across all subjects that reflects an accurate summary of full connectome effects (Bellec et al., 2015). Subjects’ T1-weighted images were segmented using SPM12 (www.fil.ion.ucl.ac.uk/spm/software/spm12/), yielding a grey matter (GM) and white matter (WM) mask in the T1 native space for each subject. The entire GM was then parcellated into 471 regions of interest (ROIs), each representing a network node by using a subparcellated version of the Harvard-Oxford Atlas (HOA), (Tzourio-Mazoyer et al., 2002), defined originally in MNI space. The T1-weighted image was then nonlinearly normalized to the ICBM152 template in MNI space using fMRIB’s Non-linear Image Registration Tool (FNIRT, FSL, www.fmrib.ox.ac.uk/fsl/). The inverse transformations were applied to the HOA atlas in the MNI space, resulting in native-T1-space GM parcellations for each subject. Then, T1-weighted images were co-registered to native diffusion space using the subjects’ unweighted diffusion image as a target; this transformation matrix was then applied to the GM parcellations above, using FSL’s FLIRT linear registration tool, resulting in a native-diffusion-space parcellation for each subject.

#### Functional connectivity

Functional connection matrices representing task-related connection strengths were estimated using a correlational psychophysical interaction (cPPI) analysis used previously by our group (Davis et al., 2017) and others (Fornito et al., 2012) to estimate a whole-brain connectivity matrix that describes task-related interactions between brain regions. Briefly, the model relies on the calculation of a PPI regressor for each region (or node), based on the product of that region’s time course and a task regressor of interest, in order to generate a term reflecting the psychophysical interaction between the seed region’s activity and the specified experimental manipulation.

The convolved task regressors from the univariate model described above were used as the psychological regressor, which were originally coded as the four Set-Size-modulated delay regressors (range = 1-4); all regressors are mean-adjusted in FSL. The correct-trial delay-period regressors were each multiplied with two network time courses for region *i* and *j.* The partial correlation *ρ_PPI_i_. PPI_J·z__* was then computed, removing the variance *z*, which includes both the psychological regressor and the time courses for regions *i* and *j*, as well as constituent noise regressors including 6 motion parameters and noise regressors coding for the concurrent signal in WhM and CSF during each run. This cPPI analysis resulted in 4 separate output matrices, comprising connectivity delineated by WM load. Task-related connectivity was estimated from the resulting output matrices; negative connections were included in these analyses, as they may inform important, explicit interpretations about how networks may be segregated (Braun et al., 2012). In order to evaluate the behavioral significance of these patterns, two additional matrices were computed based on the incorrect-trial delay-period regressors at the two highest difficulty levels; only the highest two levels were included due to insufficient number of incorrect trials at the easiest difficulty levels.

Lastly, in order to summarize system-wide behavior in the task-related network, a previously reported measure of system segregation was applied with modifications (Chan et al., 2014). This measure is calculated as the difference between the mean magnitudes of between-system correlations from the within-system correlations as a proportion of mean within-system correlation.

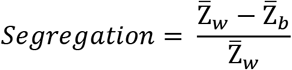

Where 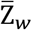 is the mean r-values between nodes of one partition, module, or system (similar to within-module degree or WMD), and 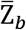 is the mean of r-values between nodes of separate partitions (similar to between-module degree or BMD, Davis et al., 2017). Accordingly, values greater than 0 reflect relatively lower between-system correlations in relation to within-system correlations (i.e., stronger integration of systems), and values less than 0 reflect higher between-system correlations relative to within-system correlations (i.e., diminished integration of systems). This segregation measure was then modified by taking *1-Segregation* to reflect system *integration,* or the inverse effect of segregation. In subsequent connectivity analyses, one older adult participant’s average network integr ation value was greater than two standard deviations below the mean, and they were thus excluded from this analysis. All plots were created using R (R Core Team, 2012) and ggplot (Wickham and Wickham, 2007). All connectivity data was visualized using BrainNet Viewer (http://www.nitrc.org/projects/bnv/).

## Results

### Behavioral results

There were no significant group differences between young and older adults in either Starting Load (t_66_ = 0.46, p = 0.65) or WM capacity (82% point of sigmoid-fitted accuracy) values (t_66_ = 1.26, p = 0.21). This suggests that both age groups were fairly equivalent in baseline performance level. Results from the linear and logistic regression models demonstrated that increasing WM Difficulty Level corresponded to lower accuracy (χ^2^ = 91.11, p < 2.2e-16) and slower RTs (X^2^ = 63.37, p = 1.7e-15). Additionally, while there was no main effect of group on accuracy, older adults demonstrated significantly slower RTs (X^2^ = 12.30, p = 4.5e-4). This lack of group difference in accuracy is not surprising, as previous studies have demonstrated that titrating for individual performance largely removes age differences in task performance that are typically found (Cappell et al., 2010; Schneider-Garces et al., 2010). Higher WM capacity also significantly predicted lower accuracy (X^2^ = 6.67, p=0.010) but not reaction time. These normalized plots show a linear trend across WM Difficulty Level (**Fig. 2A & 2B**) wherein WM load strongly predicts performance, but no group-by-difficulty interactions were found. These results suggest that age-group differences in behavioral performance can be largely mitigated by titrating task difficulty to individual ability. This titration allows brain measures to be more directly compared in young and older adults, as these differences would be unrelated to any disparity in task performance.

**Figure 2.**
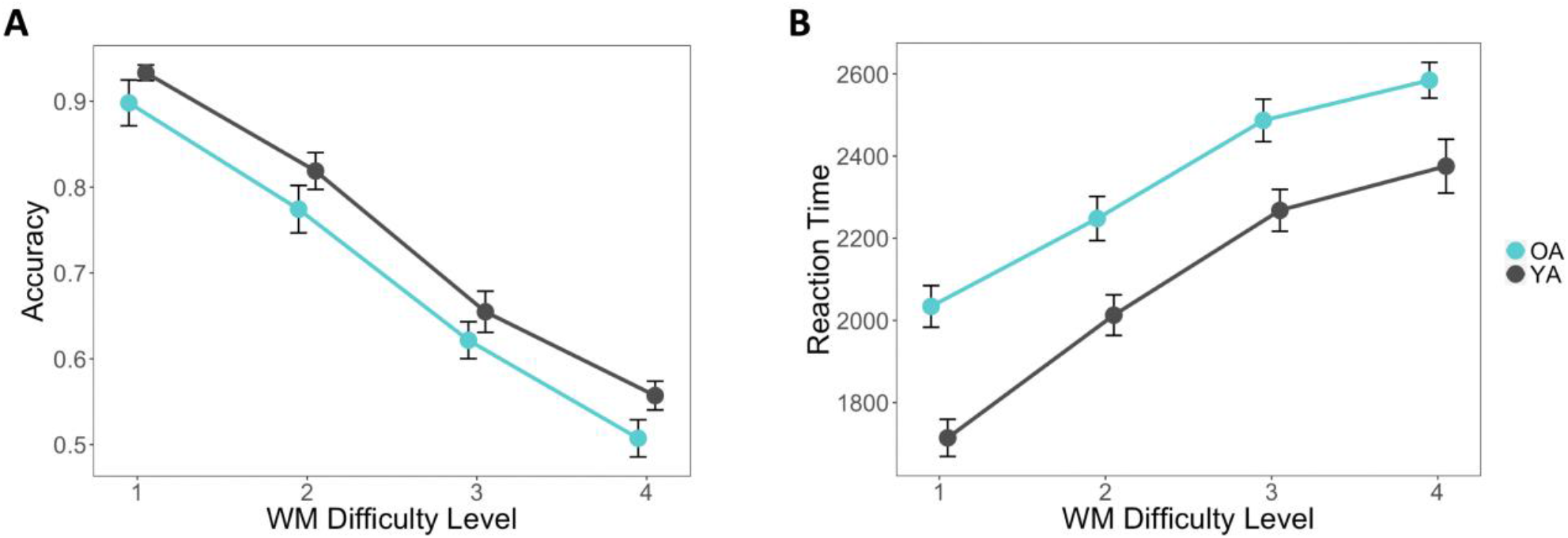
Behavioral performance in each group across difficulty, (A) showing average Accuracy ± standard error and (B) RTs ± standard error.

**Table 1.** ANOVA of factors affecting accuracy and RTsNote: X^2^ statistics and p-values were obtained by likelihood ratio tests of the full model with the effect in question against the model without the effect in question.

| Effect | Estimate | Std. Error | $\chi^2$ Value | Pr > \|t\| |
| --- | --- | --- | --- | --- |
| *Accuracy* |  |  |  |  |
| Intercept | 3.87 | 0.46 |  |  |
| Age Group | -0.32 | 0.32 | 0.95 | 0.33 |
| WM Difficulty Level | -0.81 | 0.06 | 91.11 | <2.2e-16 |
| Age x WM Difficulty | 0.03 | 0.09 | 0.13 | 0.72 |
| WM Capacity | -0.18 | 0.07 | 6.67 | 0.01 |
| *RTs* |  |  |  |  |
| Intercept | 1210.63 | 236.10 |  |  |
| Age Group | 328.68 | 90.98 | 12.30 | 4.5e-4 |
| WM Difficulty Level | 231.09 | 22.37 | 63.37 | 1.7e-15 |
| Age x WM Difficulty | -34.43 | 34.18 | 1.04 | 0.30 |
| WM Capacity | 37.71 | 39.37 | 0.89 | 0.35 |

*Note: $\chi^2$ statistics and p-values were obtained by likelihood ratio tests of the full model with the effect in question against the model without the effect in question.*

### fMRI results

As described in the introduction section, the study had three main goals: (1) investigate how age effects on WM network integration vary as a function of WM demands; (2) examine hemispheric differences in network integration in young and older adults; and (3) investigate if age-related changes in network integration relate to individual differences in WM performance. Before turning to each of these analyses, the next section describes how the task-related network was identified.

#### Task-related network identification

As illustrated by **Figure 3A**, collapsing across WM loads, the task activated the network of frontal and parietal regions typically found in fMRI studies of WM (Cabeza and Nyberg, 2000). To select ROIs for network analyses, all commonly activated ROIs across young and older adults with an average ROI-level t-value above 2.57 were included in the task network, which resulted in a 35-node network consisting of bilateral frontoparietal and sensorimotor regions (see **Fig. 3B**). As expected given the verbal nature of the task, average activity within this network was greater in the left hemisphere in young adults (F_1,36_ = 8.21, p = 0.007), but this hemispheric asymmetry activity was not observed in older adults (F_1,29_ = 1.09, p = 0.31, **Fig. 3C**). While there was no significant hemisphere by age interaction (F_1,65_ < 1), these group-level differences may indicate greater asymmetry in younger than older adults. Average univariate activity in these regions showed a weak inverse-U pattern as a function of WM load, with no differences between groups across difficulty level (**Fig. 3D**). This finding supports previous findings of load effects disappearing when normalized to individual performance (Schneider-Garces et al., 2010), and thus may be a better measure of directly comparing brain differences rather than WM capacity differences in aging. Thus, this task-based network is utilized to more effectively investigate changes in network connectivity, such as how task regions communicate with each other at network and whole-brain levels.

**Figure 3.**
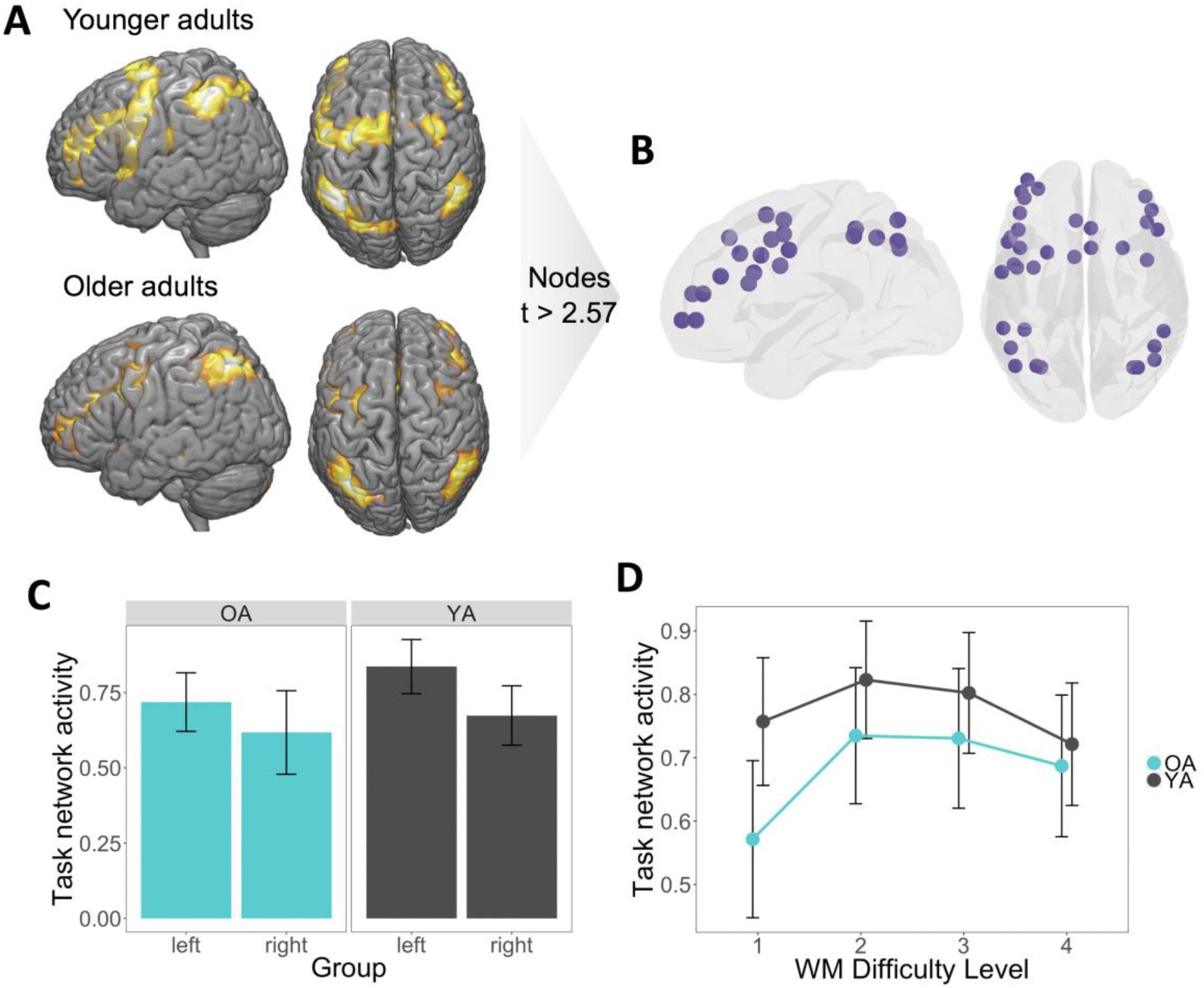
Converting univariate information into multivariate topology. **(A)** Thresholded average delay-period maps, averaging responses within each ROI in the HOA471, used to identify regions responsive to the task. **(B)** Nodes of the network with a t-value greater than 2.57 common to both groups were then assigned to the task network. **(C)** Average activity was higher in left than right hemisphere task regions. **(D)** While task network activity was higher for younger than older adults, univariate activity within the task network did not differ across WM Difficulty Level between age groups.

#### Network integration and task difficulty

The first goal of the study was to investigate how age effects on WM network integration vary as a function of WM demands. Specifically, we tested the hypothesis that *as task demands increase, the WM network becomes more integrated with the rest of the brain and this effect is greater for older than younger adults* (Hypothesis 1). As illustrated by **Figure 4**, the results were consistent with this hypothesis: as a function of WM Difficulty Level, network integration decreased slightly in young adults, but increased substantially in older adults, particularly for the hardest level (Level 4: t_65_ = −4.21, p = 8.1e-05). The interaction between group and load was significant (F_1,196_ = 5.72, p = 0.018). This finding is consistent with the univariate activity evidence for the CRUNCH hypothesis (Reuter-Lorenz and Cappell, 2008; Low et al., 2009; Cappell et al., 2010; Schneider-Garces et al., 2010), but it extends this evidence by showing that demand-related over-recruitment in older adults involves changes in network integration.

**Figure 4.**
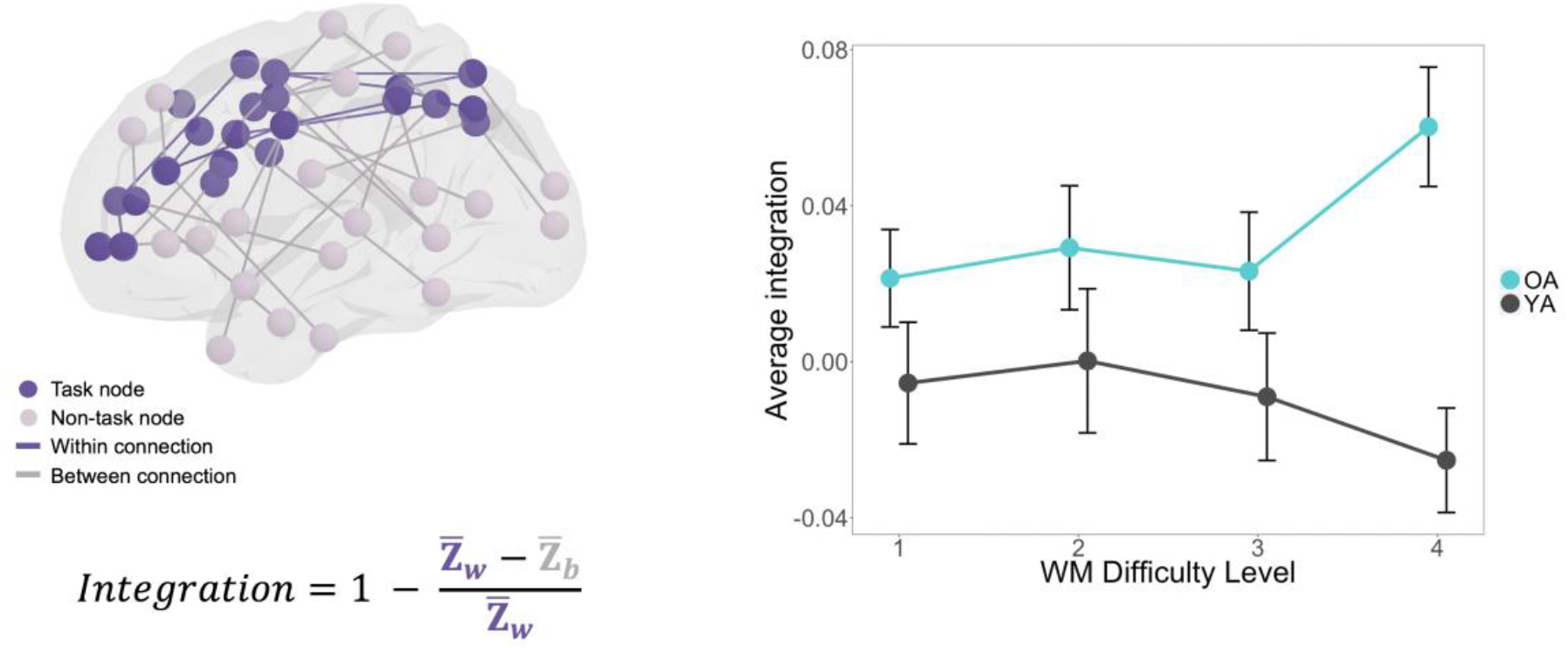
Components of integration value determination and parametric effects of WM Difficulty Level on network integration (± standard error, across WM Difficulty Level).

#### Hemispheric differences in network integration

The second goal of the study was to identify which regions are driving the integration effect by using informed decompositions of the networks examined above. First, we sought to test whether hemispheric differences in network integration in young and older adults help to explain the observed pattern above. This approach is motivated by the common finding that activations are less lateralized in older adults than younger adults. In particular, we tested the hypothesis that *during a left-lateralized verbal WM task, older adults show greater demand-related network integration in the right-than the left-hemisphere* (Hypothesis 2). To do so, integration measures were calculated separately within the left and right hemispheric components of the task network (**Fig. 5A**). Consistent with the Hypothesis 2, there was a significant difficulty by age interaction in the right hemisphere (F_1,196_ = 3.90, p = 0.048), but not in the left hemisphere (F_1,196_ = 1 68, p = 0.19; **Fig. 5B**), although there was no difficulty by age by hemisphere interaction when entered in a 3-way ANOVA. This finding suggests that age-related increases in network integration were driven by greater integration of the right hemisphere task nodes. To further investigate this effect, we split the connectivity of right-hemisphere task nodes with other right-hemisphere nodes (right-right) and with left-hemisphere nodes (right-left) focusing on the difference between Difficulty Levels 3 and 4 since these consecutive levels demonstrated the greatest interaction effect. As illustrated by **Figure 5C**, both right-right and right-left connections showed greater integration with WM demands in older adults. An ANOVA of 2 (age: young vs. old) by 2 (difficulty: Level 3 vs Level 4) by 2 (connection type: right-right, right-left) showed a significant age by difficulty interaction (F_1,192_ = 5.45, p = 0.02), but no other interactions. These results are consistent with univariate activity evidence for the HAROLD model (Cabeza and Dennis, 2013), but it extends this evidence by showing that age-related contralateral recruitment is associated with greater between-network connectivity of the contralateral hemisphere network both within- and across-hemispheres.

**Figure 5.**
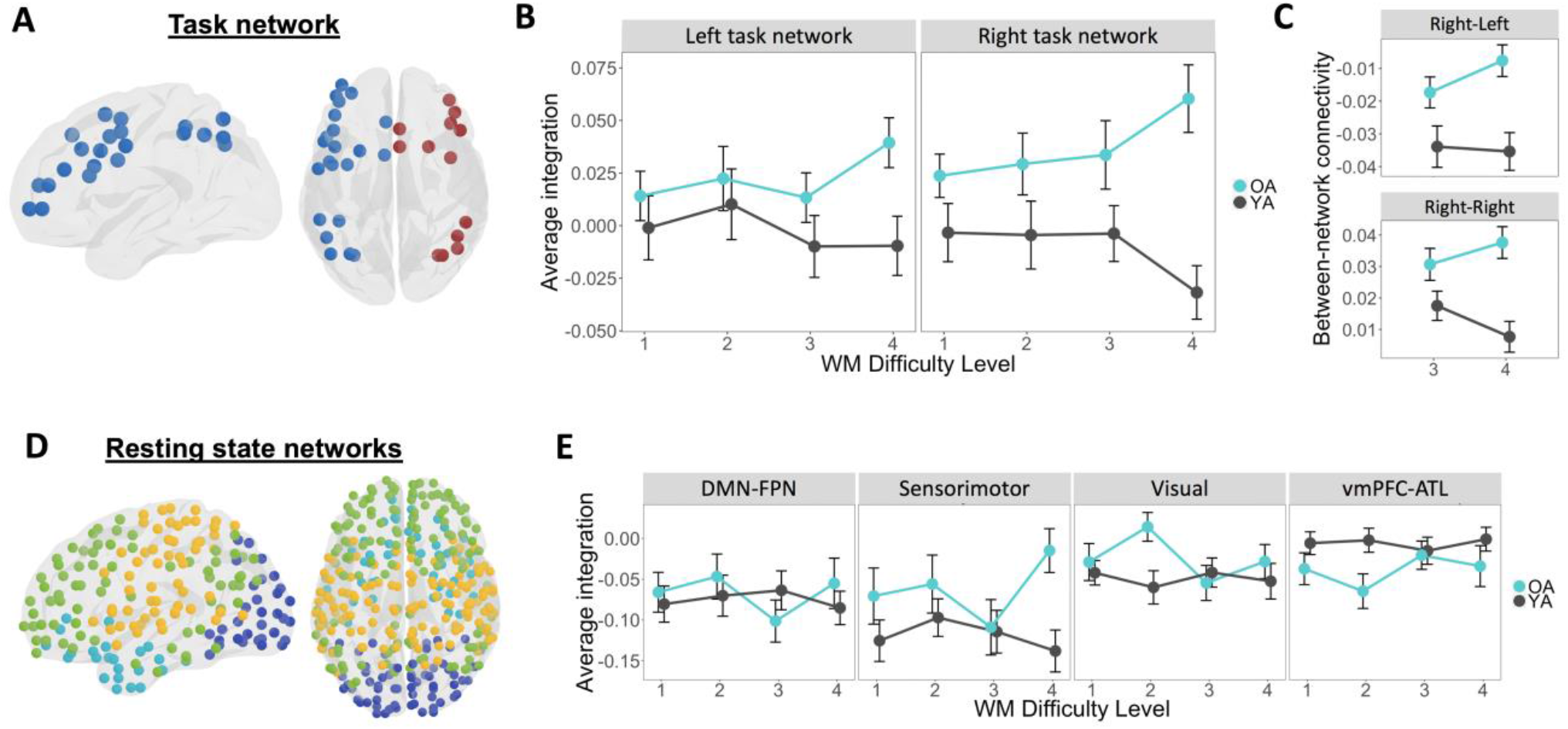
Network integration of subdivided task and resting state networks. **(A)** Parcellation of the task network split by hemisphere. **(B)** Average integration ± standard error of left-hemisphere and right-hemisphere task networks treated separately. **(C)** Right hemisphere task region between-network connectivity with right and left non-task regions. **(D)** Data-driven parcellation of modules determined by average resting state data across young and older adults. **(E)** Average integration ± standard error of each module as defined by resting state.

A potential concern of the hemispheric differences in network integration is that it could be driven by differences in canonical intrinsic networks, such as the default mode network (DMN). A growing body of evidence suggests that older adults show less modular architecture (Betzel et al., 2014), and that relationships between frontal control and DMN regions become less decoupled with age (Spreng and Turner, 2019). To investigate this issue, our second approach was to investigate the alternative hypothesis that the task-based integration was driven by changes in the modularity of resting-state networks. The average connectivity matrices of resting state were combined across young and older adults and common modules were identified. This analysis yielded four modules of relatively consistent size, labeled according to the corresponding region locations, excluding cerebellar regions: ventromedial PFC and anterior temporal lobe (vmPFC-ATL), default mode network and frontoparietal network (DMN-FPN), sensorimotor, and visual (**Fig. 5D**). The integration of the task network was computed relative to the five individual modules to determine whether the previously shown interaction in network integration was related to any standard networks (**Fig. 5E**). Integration was assessed between each of the defined modules and the whole brain, rather than the task network, using a repeated-measures ANOVA for each module. There were no significant interaction effects in any module. These results suggest that the task-network integration effect did not emerge due to increased connectivity with any specific resting state network, and that integration is perhaps best explained by a laterality decomposition within task-related regions rather than more global canonical resting state networks.

Lastly, in order to address the possibility that connectivity differences were driven by simple BOLD differences, the relationship between task-network univariate activity and integration in the task-related regions (i.e., nodes) was also investigated across all task difficulty conditions. No significant relationships between BOLD activity and Integration emerged at any level. Even in the most difficult condition, there was no trend between integration and activity in younger or older adults (YA: r = −0.04, p = 0.62; OA: r = 0.00, p = 0.99), suggesting the observed age-related increases in integration in the task network cannot be attributed to either increases or decreases in univariate activity.

#### Network integration and individual differences in WM performance

The third goal of the study was to investigate if age-related changes in network integration relate to individual differences in WM performance. Specifically, we tested the hypothesis that *age-related WM network integration would be associated with WM ability in older adults* (Hypothesis 3). As illustrated by **Figure 6A**, the results were consistent with this hypothesis: in older adults, there was a significant correlation between network integration in the most difficult condition and WM capacity (criterion score, r = 0.37, p = 0.048). In younger adults, in contrast, this relationship was not significant (r = −0.26, p = 0.18;). The correlation of WM capacity with integration was also examined at the easiest difficulty level to determine whether this effect was specific to the highest difficulty level. Neither correlation was significant (young adults: r = −0.07, p = 0.67; older adults: r = 0.09, p = 0.66; **Fig. 6B**). A Fisher r-to-z transformation was used to assess the significance of the difference between the brain-behavior correlations in younger and older adults. This difference was also significant only in the highest difficulty level (z = 2.54, p = 0.01), suggesting that network integration serves a different functional purpose in younger versus older adults. This tendency for older adults with higher WM capacity to integrate more in the difficult task condition implies that this increase in integration is an adaptive benefit to WM. This pattern of correlations also suggests that in younger adults, the segregation of the task network from non-task regions is similarly adaptive, but in the reverse direction; such a pattern of greater segregation associated with positive outcomes for behavior is consistent with other studies finding a link between increase modularity and WM performance in younger populations (Stanley et al., 2014; Braun et al., 2015).

**Figure 6.**
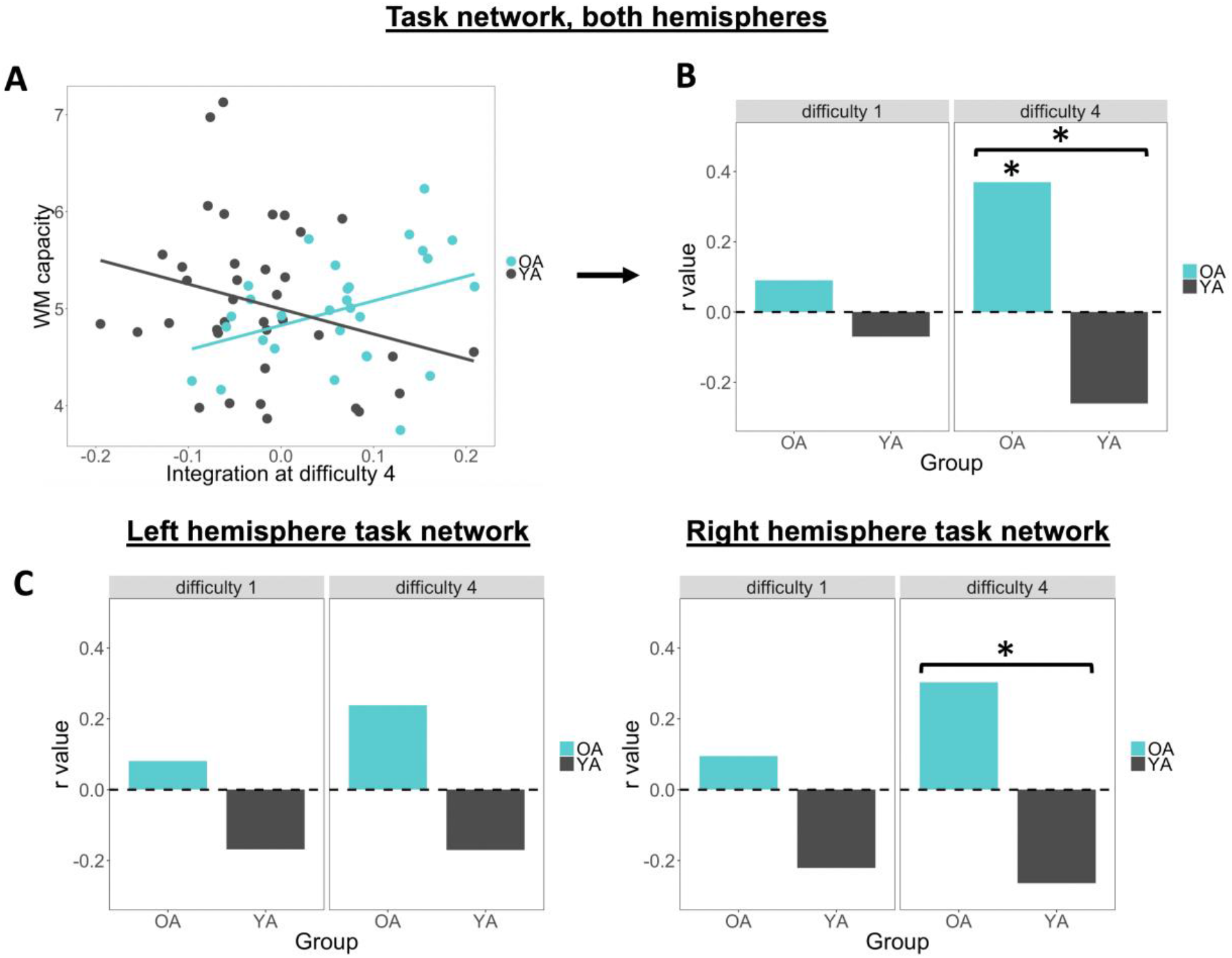
Relationship of task-network integration and WM capacity. **(A)** Task-network integration in the most difficult task condition positively correlates with individual WM capacity scores in older adults. **(B)** r values of these correlations are significant for older adults with a significant difference across group only in the high difficulty level. **(C)** Split by hemisphere, the difference in correlation values is significant only for the Right hemisphere in the high difficulty level.

To better understand hemispheric influences, we examined the same pattern of brain-behavior relationships when integration scores were split across left and right hemisphere task regions. Neither hemisphere produced a significant correlation value for either older adults (left: r = 0.24, p = 0.21; right: r = 0.30, p = 0.11) or young adults (left: r = −0.17, p = 0.31; right: r = −0.26, p = 0.11). As in the network integration findings reported above, the correlations also were not significant in the easiest difficulty level for either older adults (left: r = 0.24, p = 0.21; right: r = 0.30, p = 0.11) or young adults (left: r = – 0.17, p = 0.31; right: r = −0.26, p = 0.11). However, at the highest difficulty only, the right hemisphere showed a significant difference in correlation values in older compared to young adults (z = 2.24, p = 0.0251); the left hemisphere did not show a significant difference (z = 1.59, p = 0.11; **Fig. 6C**). This indicates that the integration effect’s association with WM capacity and age may be driven more by the right hemisphere than the left at high task difficulty, which is in line with the above finding of right hemisphere driving the age-by-difficulty differences in task network integration. Thus, an age-related difference in brain-behavior relationships is consistent with the above result that integration is good for older and bad for younger adult brains.

Correlations between average integration and accuracy during task performance at the highest difficulty level were also computed in both the bilateral task network and task network split by hemisphere; no significant relationships emerged (|r| < 0.2), suggesting that the relationship of integration with behavior may be driven more strongly by individual differences rather than task-related recruitment of a so-called compensatory mechanism.

## Discussion

The current study examined how aging affects network integration during the manipulation of information in working memory and how this integration influences individual differences in WM ability. The study yielded three main findings. First, as task difficulty increased, network integration decreased in younger, but increased in older, adults. Second, the demand-related increase in network integration observed in older adults was driven by stronger connectivity in the right hemisphere, which is less active in younger adults. Third, older adults with higher working memory capacity had significantly higher levels of task network integration in the most difficult condition, consistent with a compensation account. These three findings are discussed in separate sections below.

### Age effects on network integration as a function of task demands

The first finding of the study was that older adults showed a significant increase in network integration as a function of WM demands (load), whereas young adults showed the reverse pattern. To our knowledge, this is the first demonstration of a demand-related increase in task-related network integration in older adults, as well as the first report of a clear age by difficulty interaction in network connectivity. The age-related increase in network integration is generally consistent with studies observing load-related increases in PFC activity in older adults (Nagel et al., 2009), but it goes beyond univariate activation findings by showing that the task-related regions over-recruited by older adults are selectively integrated with the broader cortical network. This result also extends findings of age-related increases in bivariate functional connectivity (Daselaar et al., 2006; Dennis et al., 2008; St. Jacques et al., 2009) by showing the effect at the level of whole-brain networks and revealing the global context for these bivariate interactions. Finally, while the current result fits with previous evidence of age-related increases in network integration during rest (Chan et al., 2014; Chan et al., 2017), it also expands this evidence by showing a clear link between network integration and task demands.

In contrast with older adults, integration decreased in younger adults as a function of difficulty. There is currently no widespread agreement on the benefit of more widespread network community integration, with evidence showing task demands can be associated with either increased integration (Braun et al., 2015; Hearne et al., 2017) or decreased integration (Davis et al., 2018; Mattar et al., 2018), depending on task investigated. Furthermore, there is evidence that, depending on the task, greater integration can be associated with better (Braun et al., 2015; Cohen and D’Esposito, 2016) or worse (Cohen and D’Esposito, 2016) performance. In the current study, the task was the same and opposite effects of task demands on integration were seen for older versus younger adults. Given that networks tend to show less functional segregation in older adults (Betzel et al., 2014; Chan et al., 2014; Cohen and D’Esposito, 2016), one possibility is that older adults benefit more from integration than younger adults. Opposing effects of univariate activity on WM performance in older and younger adults have been previously shown (Rypma and DE’sposito, 2000; Peira et al., 2016)., suggesting that younger and older adults recruit task-sensitive regions differently in response to shifting cognitive demands.

The first finding that older adults showed increased network integration is consistent with univariate activity supporting the CRUNCH hypothesis (Reuter-Lorenz and Cappell, 2008; Low et al., 2009; Cappell et al., 2010), while also showing that faster recruitment in older adults at higher task demands is not limited to the activity of individual regions. In univariate fMRI studies, manipulating WM load typically results in an inverted-U fMRI activity pattern, first increasing with task difficulty but then declining at higher levels of difficulty (Low et al., 2009; Cappell et al., 2010; Schneider-Garces et al., 2010; Vidal-Pineiro et al., 2017). In older adults, activity increases faster at lower levels of difficulty, but it also declines faster at higher difficulty levels. These findings are postulated to result from processing inefficiencies in older adults who over-recruit neural resources at lower levels of demands and, as a result, do not have additional resources at high demand levels. We did not observe a decrease in network integration at the highest difficulty level, suggesting that network integration behaves differently than univariate activity and can keep increasing as the task becomes very difficult.

### Hemispheric differences in network integration in young and older adults

The second finding of the study was that increases in demand-related network integration in older adults were driven by connectivity changes in the right hemisphere. This finding fits with evidence for the HAROLD model (Cabeza and Dennis, 2013) showing age-related increases in univariate activity in the hemisphere less activated in younger adults, such as the right hemisphere in the current verbal WM task. Previous results speak in favor of the idea that bilateral activation patterns in the PFC are often associated with higher task demands across the lifespan (Belger and Banich, 1992; Davis and Cabeza, 2015), and therefore suggest a flexible bilateral cortical mechanism by which older adults may maintain youthful levels of performance. The current laterality finding, however, provides new insight by showing the consequence of these changes at the level of whole-brain networks. Interestingly, right-hemisphere network integration increases in older adults were mediated both by connections within the right hemisphere and by connections between the right and the left hemisphere. The latter observation is consistent with previous results showing greater bivariate connectivity between left and right DLPFC in older than younger adults (Davis et al., 2012). This therefore suggests that the bilateral compensatory pattern may not be limited to strictly contralateral regions of the brain.

In contrast with hemispheric differences in connectivity, when our connectivity patterns were decomposed by their membership to canonical resting-state networks, none of these networks showed a significant age by difficulty interaction in network integration (**Fig. 5E**), suggesting that canonical parcellations may not be sensitive to such task-related changes in network connectivity. Despite the popularity of resting state networks, it is unlikely that these partitions are representative of task-based networks (Davis et al., 2016), which could explain lack of integration in these resting state networks. Thus, while a growing body of evidence from resting-state data suggests that older adults show less modular architecture (Betzel et al., 2014), our finding highlights the importance of task specificity in the context of investigating network connectivity, as using standard network definitions may obscure effects specific to a unique task-related network.

### Network connectivity and WM capacity

The third main finding of the study was that older adults with higher WM capacity showed significantly higher network integration in the most difficult condition. While the association between higher integration and better behavioral performance has been shown in resting state data (Sala-Llonch et al., 2014), this is, to our knowledge, the first time this relationship is found in task-based functional connectivity. Interestingly, while WM capacity was found to relate to task network integration, no such pattern emerged with overall accuracy or reaction time at any given WM difficulty level. It is possible that the individual titration of difficulty may have obscured such effects, or that this finding reflects a relationship with individual ability rather than successful task performance. The lack of difference in behavioral performance between groups, thus, may be attributed to different patterns of network integration at high difficulty levels being utilized as alternative, and equally successful, processing strategies, resulting in equivalent WM capacity between age groups.

The significant difference in brain-behavior correlations between younger and older adults (**Fig. 6E**) provides some evidence for this interpretation. Given that network integration in the current study increased with task demand and was driven by the hemisphere less engaged by younger adults, the link between more widespread connectivity and WM performance provides further support for the compensatory interpretation of both CRUNCH and HAROLD (Park and Reuter-Lorenz, 2009; Cabeza and Dennis, 2013). Taken together, these findings point to the possibility that younger and older adults differ fundamentally in their approach to the WM task. The factors driving this age-related reorganization in processing are unknown, but the current study sheds some light on the topology of that reorganization. Working memory tasks, in particular, have been associated with 5Hz theta-band coupling between frontal and parietal regions during the memory retention period (Jensen and Tesche, 2002; Siegel et al., 2009) which increase parametrically with memory load. Empirical findings demonstrate that WM representations can be maintained in the absence of sustained activity during the delay in a distributed set of regions linked by oscillatory activity (LaRocque et al., 2013). The extent to which distributed activation/connectivity and distributed representation drive age-related reorganization has not been studied, but a recent analysis suggests these measures are largely decoupled (Morcom and Henson, 2018). In a recent definition of compensation, we have outlined the possibility that functional compensation in older adults reflects not simply the same cognitive process extended to new cortex, but the recruitment of additional cognitive processes not utilized in younger adults (Cabeza et al., 2018). Thus, the finding that successful performance in older adults relies on greater integration suggests that this group may rely on a more distributed set of cues (or a more flexible cognitive strategy) to perform the same task as younger adults.

### Conclusions

In sum, the study yielded three main findings. First, as task difficulty increased, younger adults showed decreased network integration, whereas older adults showed increased network integration.

Second, age-related increase in network integration was driven by increases in right hemispheric connectivity to both left and right cortical regions. Lastly, older adults with higher working memory capacity demonstrated significantly higher levels of network integration in the most difficult condition. The findings are generally consistent with two popular theories regarding age effects on brain function, CRUNCH and HAROLD, as well as with the compensatory interpretation of these effects, while also extending the evidence for these theories from univariate activity to network architecture.

## Acknowledgments

This research was funded by grant support from the National Institute of Aging grant # U01AG050618, and in part by the National Institute of Health Intramural Research program (ZIAMH00295). The authors wish to thank Duy Nguyen and Wesley Lim for their assistance with running experimental sessions and data processing. This study was based on a study pre-registered on ClinicalTrials.gov (NCT02767323).

